# Clonal reconstruction from co-occurrence of vector integration sites allows accurate quantification of expanding clones in vivo

**DOI:** 10.1101/2021.08.10.455805

**Authors:** Sebastian Wagner, Christoph Baldow, Andrea Calabria, Laura Rudilosso, Pierangela Gallina, Eugenio Montini, Daniela Cesana, Ingmar Glauche

## Abstract

High transduction rates of viral vectors in gene therapies (GT) and experimental hematopoiesis ensure a high frequency of gene delivery, although multiple integration events can occur in the same cell. Therefore, tracing of integration sites (IS) leads to misquantification of the true clonal spectrum and limits safety considerations in GT.

Hence, we use correlations between repeated measurements of IS abundances to estimate their mutual similarity and identify clusters of co-occurring IS, for which we assume a clonal origin. We evaluate the performance, robustness and specificity of our methodology using clonal simulations. The reconstruction methods, implemented and provided as an R-package, are further applied to experimental clonal mixes and to a preclinical model of hematopoietic GT.

Our results demonstrate that clonal reconstruction from IS data allows to overcome systematic biases in the clonal quantification as an essential prerequisite for the assessment of safety and long-term efficacy of GT involving integrative vectors.

## Introduction

Gene therapy (GT) approaches aim to compensate the missing functionality of a mutated gene by the insertion of one or more corrected copies of the same gene into the genome of patients’ cells. Integrative viral vectors, such as gamma retro viral (gRV) and lentiviral vectors (LV), are clinically important vehicles to realize the permanent integration of a therapeutic transgene into the genome of hematopoietic stem and progenitors cells (HSPC) to establish the expression of the defective gene in all the cells of the hematopoietic systems (Ferrari, Thrasher, & Aiuti, 2021). The semi-random integration site (IS) of the target sequence within the host genome represents an inheritable “fingerprint” allowing to track the progeny of the targeted cells over time and in different locations. However, the permanent gene transfer confers the risk of disturbing a cell’s genetic program potentially leading to uncontrolled proliferation (Cavazzana, Bushman, Miccio, Andre-Schmutz, & Six, 2019). Since the first reports about insertional mutagenesis events in early clinical GT trials (Hacein-Bey-Abina et al., 2003), the continuous monitoring of IS abundances has become a standard method to detect aberrant and potentially malignant clonal expansion (Cavazzana-Calvo et al., 2010). Therefore, corresponding protocols for the assessment of safety and efficacy are routinely implemented in GT trials.

The availability of new molecular approaches for IS retrieval (Firouzi et al., 2014) utilizing next generation sequencing (NGS) has greatly improved the efficiency to identify and quantify the abundance of multiple IS within one sample (Beard, Adair, Trobridge, & Kiem, 2014). Although those methods are steadily improved to achieve a better correlation between input material and the number of identified IS, there is still a number of technical challenges. Nevertheless, it is generally accepted that the IS abundance also corresponds to the clonal abundance, although, this interpretation is only valid if each clone is solely marked by a single IS (Berry et al., 2012). Aiming towards high transduction rates among initially transplanted cells, target cells commonly integrate more than one virally transduced genetic sequence. The incorporation of multiple IS within individual cells strongly affects the interpretation of quantitative IS analysis. For the example in Figure 1, we consider three clones with two, four or six IS. While the IS time series in Figure 1A appears balanced, correct assignment of the IS to their corresponding clones and averaging IS abundance per clone leads to a different interpretation in Figure 1B, in which one clone clearly dominates. Unfortunately, the missing association between IS and clones is a fundamental challenge in GT applications that increases the risk of overestimating the overall clonality and underestimating single clonal abundances. Especially in clinical settings, in which the safety assessment of GT trials is explicitly based on relative IS abundances (Aiuti et al., 2013; Biffi et al., 2013; Hacein-Bey Abina et al., 2015; Kohn et al., 2020; Marktel et al., 2019; Sessa et al., 2016), it is essential to know about and correct for multiple integrations.

**Figure 1.**
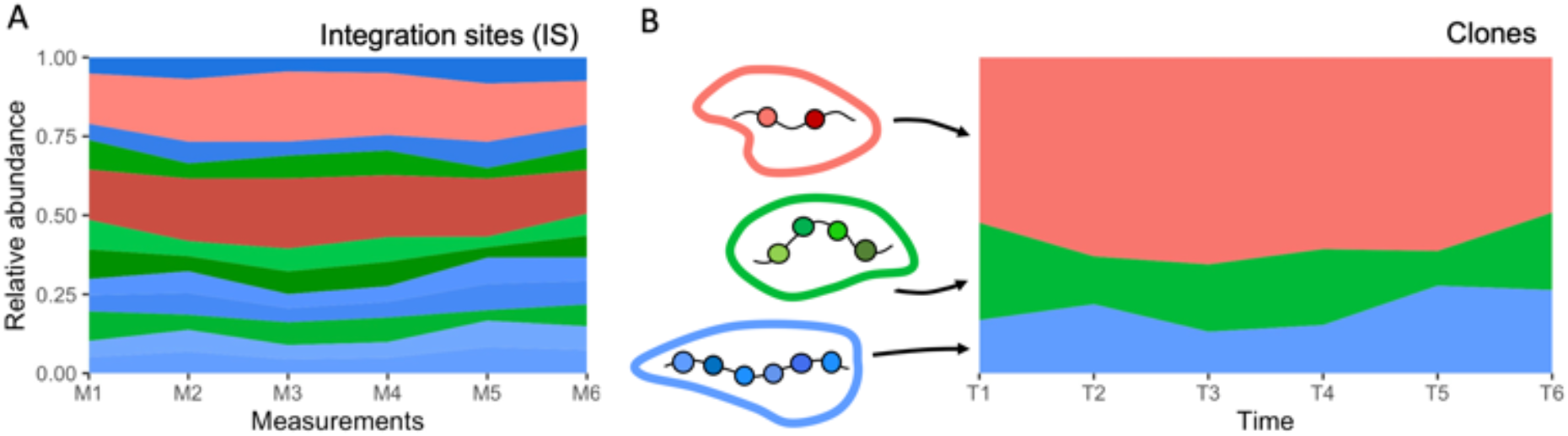
Sketched example of an IS based time series resulting from the contribution of three clones. **A** stable detection of 12 IS over different measurements (relative abundance). Although the colors indicate the clonal origin, this information is usually not known *a priori*. **B** reconstruction of the common clonal origin of the different IS allows to recalculate the true, underlying clonal time course.

There are currently no methods available to address this issue and to correct clone size quantification. Therefore, we developed a novel bioinformatic approach to detect co-occurrent IS within the same clone and provide an R software package *MultIS* for the corresponding analysis. Our approach is based on the idea that two IS of the same clone appear in a constant relative frequency to each other while IS from different clones will change their mutual relative frequency according to the corresponding different clone sizes. We use a mathematical modeling approach to illustrate how the identification of mutual correlations between all pairs of IS can be used to identify sets of IS with high similarity, suggesting the same clonal origin. We particularly employ mathematical modeling to demonstrate both the potential and limitation of this approach. We further validate our method using *in vitro* data obtained by mixing multiple cell clones with different but known IS. Finally, we apply our pipeline to a mouse model that mimics a real case scenario of HSPC-GT. Hence, we developed and experimentally validated a new method that accurately identifies source clones from observed IS, improving downstream analyses for preclinical and clinical studies.

## Results

### Clonal reconstruction for simulated time series data

We developed a bioinformatic pipeline to identify IS that belong to the same clone. Our suggested method is based on the idea that multiple IS of the same clone appear with similar relative abundances. If the source clone is small, all its IS should present with low abundance, while IS of a dominant clone should all appear more frequent. Following an initial *filtering* step we calculate pairwise *similarities* among all IS which are fed into a *clustering* algorithm that identifies co-occurring, *clonaly* related IS (Figure 2).

**Figure 2.**
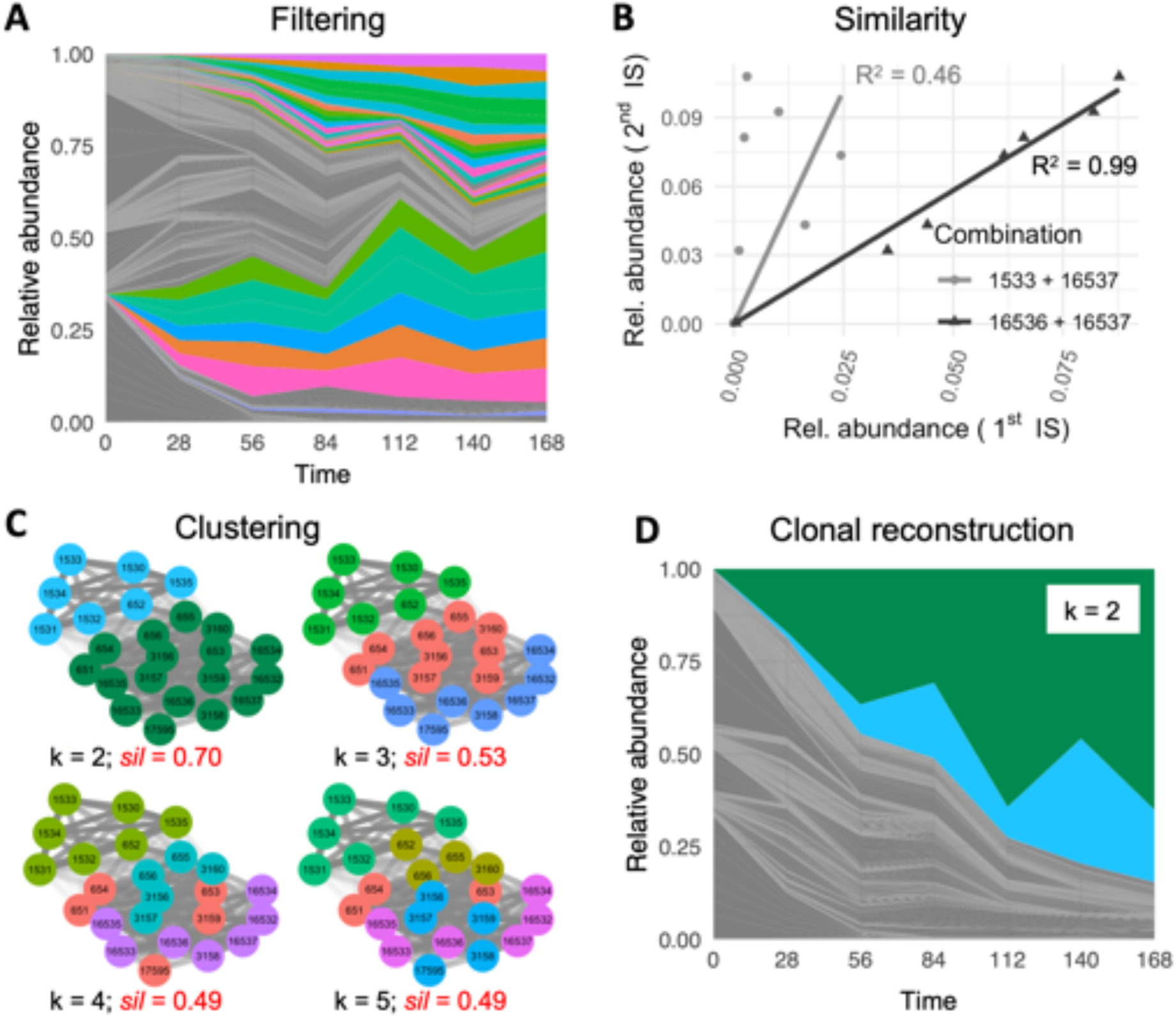
Overview of the reconstruction pipeline. **A** example of a time series of IS abundance after applying a filtering step (IS > 1% at the last time point). Grey colors indicates IS that did not pass the filtering. **B** two examples for the calculation of the similarity score R^2^ between two pairs of IS. The smaller residuals between triangles and the black line indicate a higher similarity between IS 16536 and 16537 (R^2^ = 0.99) as compared to IS 1533 and 16537 (R^2^ = 0.46), which are shown in grey and present with larger residuals. **C** displays network representations of the corresponding similarity matrix *S* in which the shading of the edges indicates the mutual correlation between the IS. We show four different optimal clusterings for increasing values of required clusters k = 2 to 5, for which the coloring of the nodes represents the obtained clustering. Using the silhouette score *sil* (provided in red) we compare the overall clustering quality between these results, indicating that for the particular data set, k = 2 clusters optimally reflects the structure of the similarity matrix *S*. **D** displays the clonal time series which has been reconstructed by assigning IS of one cluster to the same clone. IS that did not pass the filtering, are still shown in grey.

In order to demonstrate the general feasibility of our approach and to learn about its limitation, we implemented a scalable mathematical model mimicking time series of clonal development. In brief, our model describes the proliferation and the differentiation of a stem cell population. Implementing this approach as a stochastic, single cell-based model allows to follow the progeny of each individual cell and thereby to track clonal developments (Baldow et al., 2016). Depending on the choice of parameters we can influence the heterogeneity of the initialized cell clones with respect to their tendency for differentiation (encoded by the standard deviation of the mean differentiation rate *δ*), and thereby regulate how fast the system will converge towards clonal dominance (Figure 3 A-C). On top of those clonal time series we superimpose several IS per clone. Technically, we sample the number of IS per clone from a Poisson distribution with mean λ which reflects the average VCN. Examples for this superposition for λ = 5 are shown in Figure 3 D-F.

**Figure 3.**
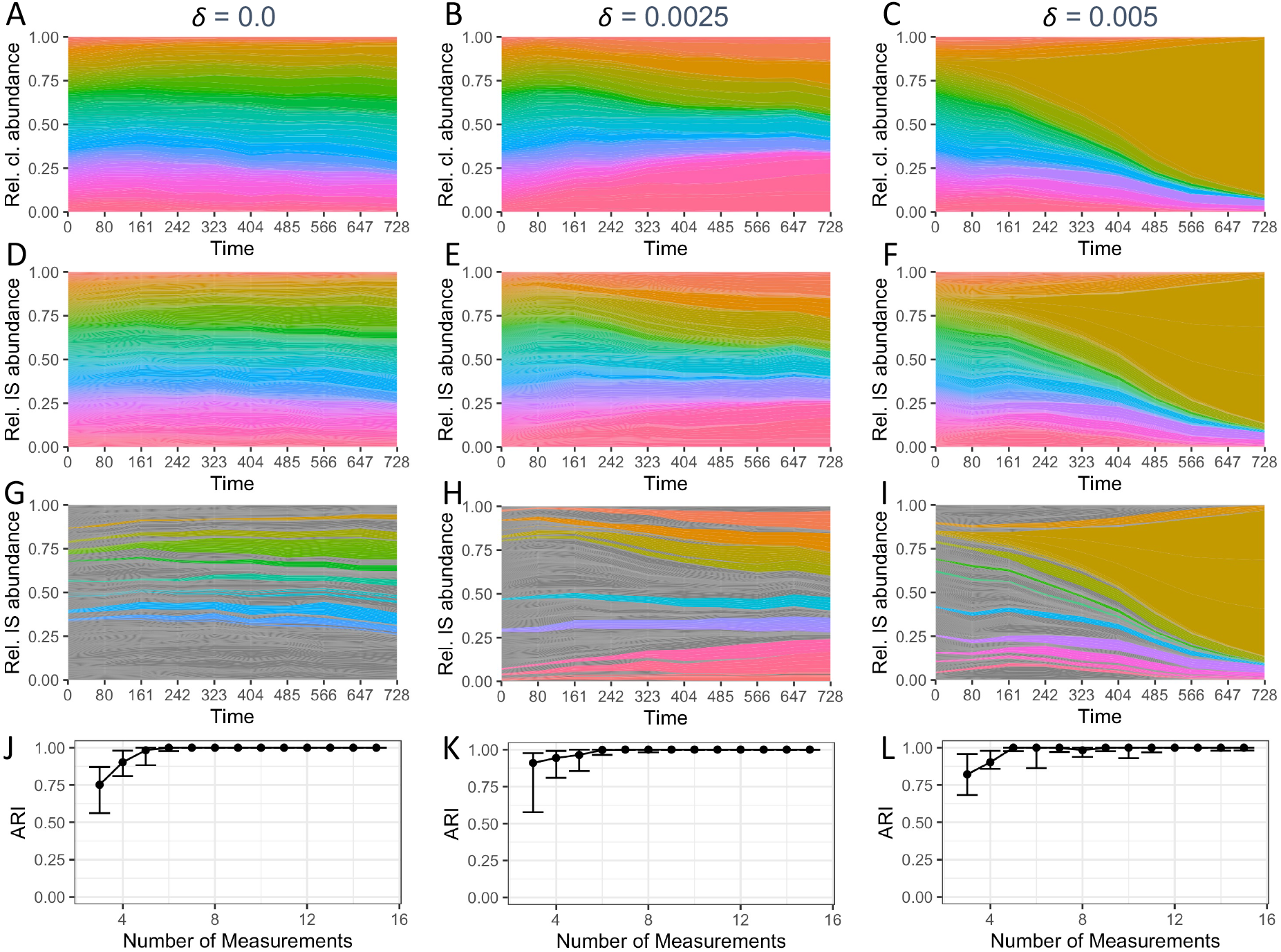
Time series of clonal development and their reconstruction based on IS measurements. **A,B,C** clonal time series for increasing values of the SD of the differentiation rate *δ* with an increasing tendency for clonal conversion. **D,E,F** corresponding time series for IS, which are superimposed in each clone. **G,H,I** remaining time series after filtering for the 5u largest IS at the final time point. **J,K,L** Quality of the clonal reconstruction process as a function of the number of available time points for IS measurements. Reconstruction quality is measured by calculating the ARI between the known ground truth and the reconstructed clustering (median ARI → 1 indicating perfect reconstruction; quantitative analysis is based on 20 independent simulations each; points indicate the median; whiskers correspond to the first and third quartile).

From these simulated clonal developments, we obtain time series of the relative IS abundance 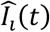 which are scalable for their clonal heterogeneity (*δ*), the average VCN λ, but also for the number of measurement time points and the level of measurement noise *σ*. It is the central advantage of the simulation model, that the ground truth, i.e. the true assignment of IS to clones is intrinsically known and can be used as a benchmark to evaluate the performance of the reconstruction process.

The bioinformatic reconstruction is solely based on the IS time series 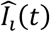 (see Materials and Methods). Briefly, the reconstruction pipeline is initiated by a filtering step to minimize the detection of spurious correlations and to focus on the clones with higher abundance. Herein, we are only considering the 50 most abundant IS at the final time point of the analysis (Figure 3 G-I). As the second step, we calculated a similarity score for any pairs of filtered IS, which is subsequently used to identify clusters of IS with highly correlated behaviors. Those clusters are interpreted as single clones with potentially multiple IS. In a final step, we recalculated the clonal abundances as an average of the IS abundance per clone and for each measurement (Supplementary Materials, Supplementary Figures S1 and S2).

We quantify the reconstruction quality using an adjusted Rand index (ARI), which compares the true association of IS to clones with the corresponding assignment obtained from the reconstruction pipeline. Figure 3 J-K indicates the median of the reconstruction quality increases as a function of the number of equally spaced measurement time points, suggesting that a reliable reconstruction can be obtained for most values of the clonal heterogeneity *δ* given that at least five independent measurements are available.

Delineating the individual clonal time series according to their assigned clusters indicates that the IS in the same cluster indeed show highly correlated behavior (Supplementary Figures S1 and S2). Using this assignment of IS to clones, a reconstructed clonal time series can be obtained which mimics the general behavior already known as the ground truth. Minor IS that did not pass the filtering step were corrected for the average number of IS per clone and are indicated as background.

We further analyzed the influence of the average VCN *λ* and the level of measurement noise *σ* on the reconstruction quality (Figure 4A-I). We observe that the reconstruction algorithm performs worse for smaller VCN, especially if the average value is as low as *λ* = 2 or smaller. In those cases, the majority of clones harbors one IS. As the clustering approach has a tendency to combine weakly correlated IS into one clone, the overall reconstruction quality is diminished. For increased average VCN *λ* a high reconstruction quality is achieved while for very high values (*λ* > 10) too few clones remain after the filtering. In an orthogonal dimension, we study the influence of noise *σ* for the measurement of each individual IS. Figure 4A-I indicates a robust reconstruction even for increasing levels of measurements noise *σ*. However, the reconstructions fails for large values of *σ* as true correlations between IS can no longer be detected.

**Figure 4.**
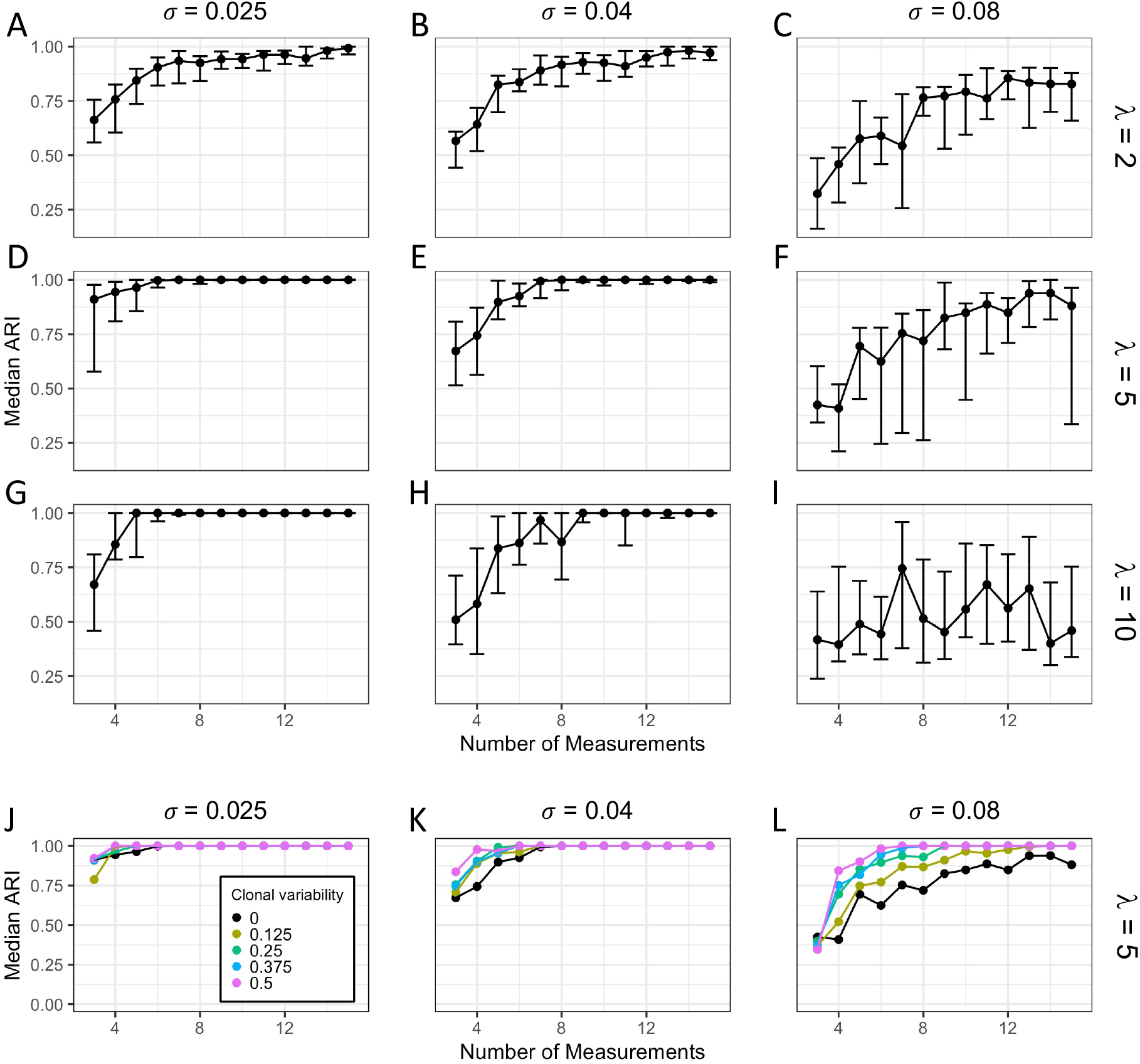
Dependencies of the reconstruction quality. **A-I** Reconstruction quality with respect to varying VCN and levels of measurement noise *σ* for standard deviation of the differentiation rate δ = 0.0025. Rows refer to the average VCN λ = 2 (**A,B,C**), λ = 5 (**D,E,F**) and λ = 10 (**G,H,I**). The column represents increasing levels of the measurement noise *σ* = 0.025 (**A,D,G**), *σ* = 0.04 (**B,E,H**) and *σ* = 0.08 (**C,F,I**). Each data point is based on 20 independent simulation runs (points indicate the median ARI; whiskers correspond to the first and third quartile). **J-L** Reconstruction quality depends on clonal variability v. Median ARI is shown as a function of the number of measurements for increasing levels of measurement noise *σ* = 0.025 (**J**), *σ* = 0.04 (**K**) and *σ* = 0.08 (**L**). The coloring corresponds to different levels of the clonal variability v, which approximates the diversity of available compartments. Each data point is based on 20 independent simulation runs.

Our results indicate that a correlation-based assessment of IS abundances is indeed suited to identify multiple IS co-occurring in the same clone.

### Assessment of IS abundance in different hematopoietic lineages improves the reconstruction process

Various studies of the hematopoietic system have shown that HSPC clones obey the tendency to preferentially contribute to one or another hematopoietic lineage (Notta et al., 2016; Scala & Aiuti, 2019; Scala et al., 2018). For example, some clones preferentially differentiate to T cells while other clones contribute more to granulocytes. For the first case we would expect to see a prominent contribution of all the clonal IS to the T cells, while the same IS should rarely be seen for the granulocytes. As the analysis of correlations between different IS benefits from varying clone sizes, we hypothesize that the assessment of IS abundance in different hematopoietic lineages can further improve the reconstruction process. In this notion, measurements of the IS abundance in different lineages are considered as independent samples closely similar to measurements at different time points.

To account for the fact that clones do not equally contribute to different lineages, we introduced an artificial clonal shift in the model simulation, quantified by parameter ν. For ν = 0, the final clone sizes correspond to the clonal time courses (i.e. 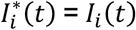), while for values of ν > 0 clone sizes for each measurement point are randomly and moderately varied, thereby affecting the abundance of all IS for each particular clone.

Clonal reconstruction based on the resulting model simulation for ν > 0 confirms our hypothesis. Figure 4J-L indicates that especially for higher values of the measurement noise *σ* there is a consistent improvement in the reconstruction quality for increasing values of the clonal variability ν. This additional level of variability, which compares to clonal measurements in distinct hematopoietic compartments, appears as a key factor to strengthen the identification of mutually correlated IS and their clonal origin. We explicitly point out, that those measurements do not necessarily require temporal separation, but can be achieved by subfractioning primary samples according to lineage identity prior to sequencing.

We conclude that IS measurements in different hematopoietic sub-compartments improve the clonal reconstruction process.

### *In vitro* validation assays confirm the validity of clonal reconstruction

We further validated our reconstruction method using an *in vitro* experimental assay. To this end we used four K562 cell clones having different and known IS: ID#27 (1 IS), ID#30 (4 IS), ID#37 (6 IS), ID#46 (10 IS). The genomic position of each clone specific IS was previously identified by SLiM_PCR (Supplementary Table S2). In order to replicate a potential *in vivo* situation and to challenge our clonal reconstruction model, we designed an *in vitro* assay where the four cell clones were mixed at different ratios (Supplementary Table S1), such that each clone specific IS was present at a predefined level of abundances. A second cell line transduced in bulk with a SINLV and having an average VCN = 1.8 was added to all these mixes to generate the background signal of small clones that could be present in a real case scenario and interfere with the detection of emerging clones. True clone abundances were confirmed using a droplet digital PCR assay.

Figure 5A illustrates the measured relative abundance of the prominent IS for the seven different mixes (left bar with black contour). Spurious IS derived from the transduced background (ranging from 7% to 60% for the different mixes) were already removed by filtering for IS > 1% in any sample.

**Figure 5.**
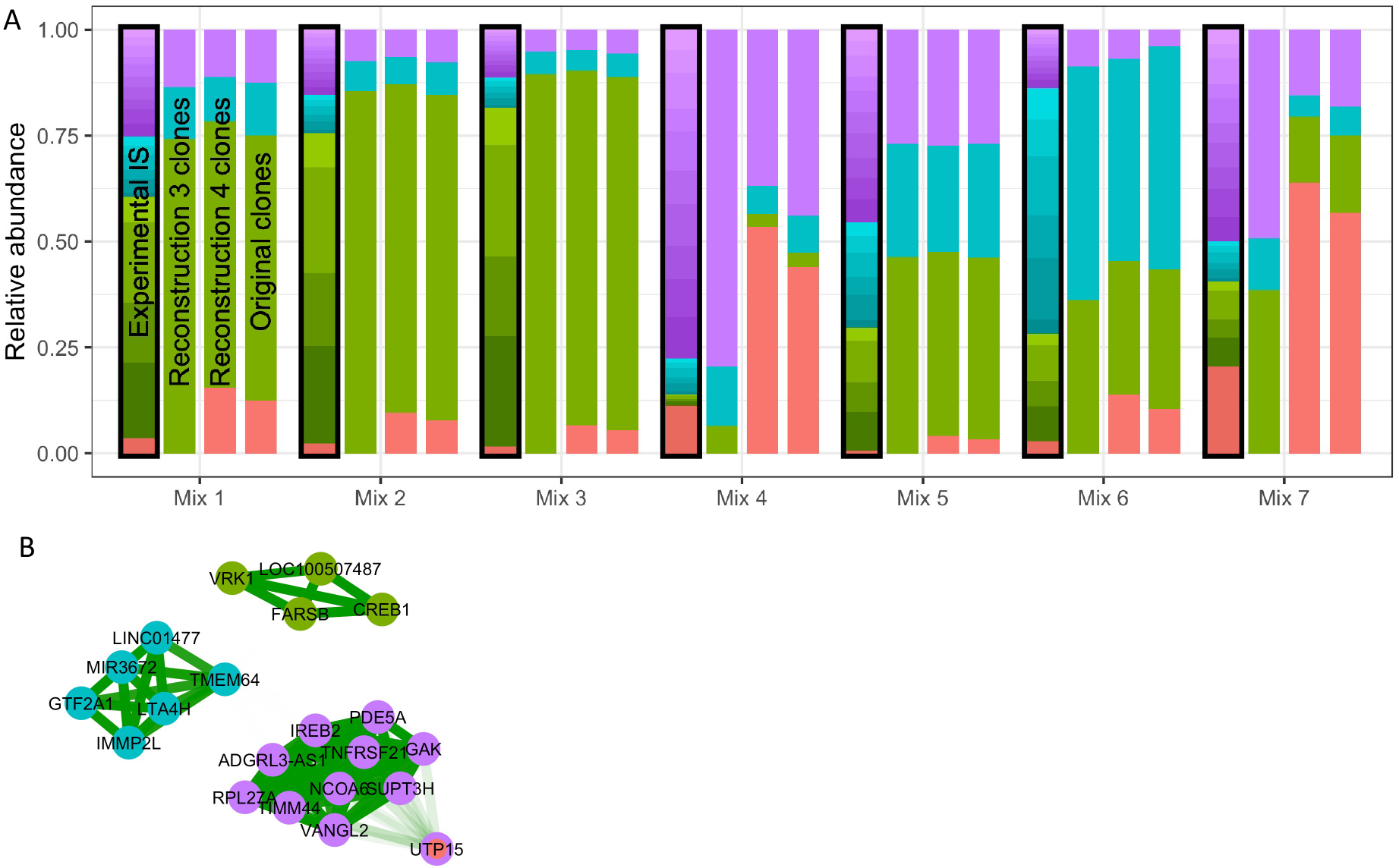
Analysis of in vitro mixes of the validation assay. **A** For each of the seven analyzed samples (mixes 1 to 7) four stacked bar plots display from left to right: the relative proportion of measured IS after filtering (black contour, highlighting the available data in an experimental/clinical context, although the assignment to one of the four clones (i.e. the coloring) is generally unknown), the relative proportion of the different clones after applying the reconstruction routine, the relative proportion of the different clones after manual correction (see text), and the relative proportion of the four original clones. **B** Network map indicating the clustering of mutually similar IS. Shading of the edges indicates the mutual similarity R^2^. The inner color of each node represent the true assignment to one of the four clones, while the outer ring corresponds to the result of the reconstruction process.

By applying our suggested reconstruction pipeline, we obtained clusters of IS which appear correlated and which are interpreted as *clones* (Figure 5A, second bar). Figure 5B illustrates the strong inner-cluster similarity (dark green edges), which allows us to correctly identify the IS belonging to three of the four clones. The example also illustrates that clone ID#27 (which is only characterized by a single IS) is falsely assigned to one of the three other clones. It appears as an intrinsic limitation of the clustering approaches that singular IS are preferentially joined to one of the other clusters. Although heuristic methods (such as the detection of bimodal similarity scores within clusters) can be implemented to detect weakly assigned IS, we recommend a prior visual inspection to identify and compensate this shortcoming. Correcting for the obvious misclassification in the given case, the *in vitro* assay confirms that the estimated clonal abundance obtained from the reconstruction pipeline (Figure 5A, third bar) closely recapitulates the respective ground truth (Figure 5A, right bar, Supplementary Table S3). The resulting approximation of the true clonal mixture outperforms a sole assessment of the IS which largely overestimates the true number of dominant clones.

Based on these results, we concluded that our method is indeed suited to reconstruct clones from IS using their abundances over different observations.

### In vivo testing on a mouse experiment

To further confirm the potential of our reconstruction method for the detection of expanding cell clones harboring multiple IS, we took advantage of a preclinical model of HSC gene therapy (GT) based on tumor prone Cdkn2a^-/-^ Lineage-(Lin-) cells. In this model, lethally irradiated WT mice are transplanted with Cdkn2a^-/-^ Lin-cells transduced at multiple copies by a neutral self-inactivating lentivirus expressing GFP (SINLV.PGK.GFP) (Cesana et al., 2014). Due to the tumor prone background of the Lin-cells, the transplanted animals develop hematopoietic malignancies with a specific kinetic (Montini et al., 2006).

We obtain measurements of IS abundance at four to five different time points in different blood lineages (B-lymphocytes (CD19+), T-lymphocytes (CD3+) and myeloid cells (CD11b+)) starting at week 4 post transplantation. Those are complemented by tissue samples obtained at autopsy, in which bone marrow, blood, spleen, thymus, and lymph nodes were collected for IS site retrieval. Following the same reconstruction pipeline as for the simulation scenario, we started off by filtering the most abundant clones with a relative abundance of more than 1% in the final blood or tissue samples (Figure 6A). Analyzing the mutual similarities between those dominant IS, we identified IS with closely correlated behavior, which appear as strongly connected clusters in corresponding network maps (Figure 6B). We interpret this pronounced, visual separation of clusters as a strong indicator of common clonal origin. For the particular example, our analysis suggests that 53 filtered IS can be optimally mapped to eight different clones. Representations of the clustered time series confirm the high inner-cluster similarity (Figure 6C).

**Figure 6.**
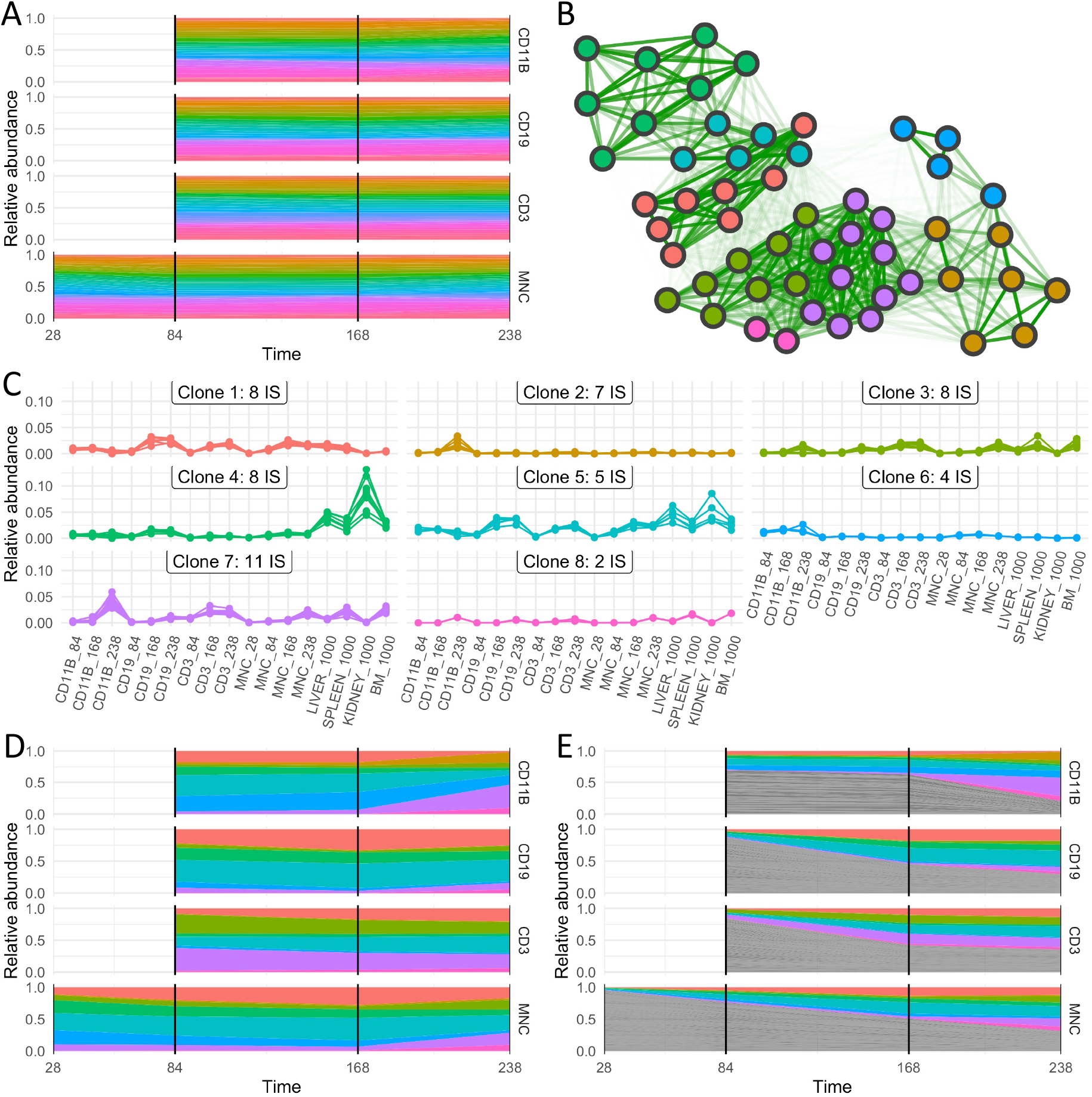
Experimental data and clonal reconstruction for wt mouse E4C. **A** relative abundances of IS as a function of time for subsets of CD11b, CD19, CD3, and mononuclear cells, for which multiple measurements are available. **B** The network map depicts the similarity between each pair of integration sites (indicated by edge brightness) superimposed by the optimal clustering obtained from the reconstruction pipeline (indicated by color of the nodes). **C** shows the time series of all IS assigned to the same clusters (preserving the color scheme). **D** corrected clonal time series for the four identified clones. **E** corrected clonal time series together with the IS that did not pass the initial filtering step (indicated in grey).

Based on this assignment of IS to a much smaller number of clones we can now translate the IS time series into a corresponding clonal time series. Figure 6D indicates a less polyclonal pattern as suggested by just accounting for the IS. Figure 6E further acknowledges less abundant IS that did not pass the initial filtering step (indicated in grey), leading to a quantitative description of dominating *clonal* dynamics over the full experimental period. Two further examples from the same experimental setting are provided in Supplementary Figures S3 and S4.

Our analysis of an experimental GT setting with known integration of multiple vectors per cell clone illustrates the applicability of our method in an application-relevant context.

### R implementation: the *MultIS* package at CRAN

In order to facilitate the application and further development of our analysis pipeline we provide a corresponding R package named *MultIS* via CRAN (https://cran.r-project.org/package=MultIS). Starting out from a list of IS abundances *I*(*t*) for multiple measurements, the package combines all individual parts of the analysis pipeline along with graphical representations. We provide a corresponding R-script file to reproduce our analysis (https://gitlab.com/imb-dev/clonal-reconstruction-figures).

## Discussion

The unbiased assessment of temporal clonal contributions to different hematopoietic compartments is limited by the co-occurrence of several IS within the same clone which can hardly be resolved experimentally. In this work we propose a novel bioinformatic pipeline that overcomes this limitation and leverages intrinsic correlations between IS abundances derived from the same clone to recover the true clonal structure. Our approach relies on the idea that IS from the same clone appear in the same relative frequency across time series measurements and among different hematopoietic cell types, while this correlation is missing for IS resulting from different clones. This concept is translated into a corresponding analysis pipeline, which we provide as a publicly available R package *MultIS*.

We first optimized and validated our method using mathematical simulations of clonal dynamics. We demonstrated that the method is broadly applicable for settings with VCN ≥ 2 in the dominating clones. Systematic variation of central model parameters identified fewer measurements and increasing measurement noise as limiting factors of the reconstruction process, as pairwise correlations of IS abundance are harder to detect. The decline in reconstruction quality can be compensated by considering different hematopoietic subcompartments with varying clonal contributions, even if they are measured at the same time point. Our simulation studies confirmed that the central source of information for the reconstruction process is based on the variability in IS abundances re-captured at different measurements. Intuitively, if all clones are always present with a highly reproducible proportion, also the IS of these clones are more or less constant over repeated measures. However, a reliable correlation can only be obtained if IS of the same clone vary in a synchronized manner, namely due to their changing clonal abundance. This observation indicates that not only temporally separated measurements but also IS analyses in different hematopoietic lineages with distinct clonal contributions at the same time point are extremely valuable to reconstruct the IS affiliation with good precision.

Applying the reconstruction pipeline to a set of *in vitro* assays in which cell clones with different numberS of IS are mixed in predefined ratios confirms the feasibility of the approach. Solely based on the quantification of IS abundance we can identify which of the IS belongs to which clone. These experiments also point towards a limitation of the automated clustering approach, as it has a tendency to misinterpret clones with unique IS. The clustering has a higher tendency to join such IS to other clusters instead of labeling them as unique. Wrong assignment of a few unique IS to identified clusters will only moderately effect the overall interpretation of the clonal reconstruction, although we strongly recommend a visual inspection of the clustering results to identify obvious misclassifications. Threshold based corrections (using e.g. the detection of a bimodal distribution of similarity score withon one cluster) can heuristically target this problem, while critical cases might need to be resolved by an experimental validation based on the sequencing of single cell colonies.

In the final step of our analysis we use the reconstruction pipeline to estimate clonal expansion in an *in vivo* model for malignant hematopoiesis. Our results confirm the hypothesis from the simulation results, namely that the assessment of different hematopoietic lineages improves the clonal reconstruction process. Although the true assignment of IS to clones is unknown for these experiments, the overall similarity and consistency of the clustered time series reinforces our primary intention that the reconstruction process can also be applied to *in vivo* settings, and transferred to GT applications.

Current studies of clonal dynamics only quantify IS abundances relative to the total observed IS and interpret those time series as independent clones. This approach can clearly lead to an overestimation of the number of clones and an underestimation of their relative contribution. However, the quantitative assessment of clonal behaviors is crucial for the evaluation of the safety of GT as well as for the interpretation of experimental studies to understand physiological hematopoiesis and related malignancies or to study hematopoietic reconstitution *in vivo*. The availability of repeated IS measurements at different time points as well as in different hematopoietic cell types, especially in a clinical context, represents the optimal data basis necessary for the successful application our suggested methodology and to reach a more thorough understanding of temporal clonal developments.

## Materials and Methods

### Reconstruction pipeline

The reconstruction pipeline provides a method to identify clones based on the tracking of IS over time and/or in different hematopoietic compartments. An initial *filtering* step (Figure 2A) restricts the analysis to prevent potential biases from under-represented clones. For the filtering of the simulated time series, we obtain interpretable results over a broad range of parameter settings if we consistently consider the 50 largest remaining IS at the final time point. For the biological data we filter IS with an abundance above 1% at the final measurements to account for the detection thresholds of the quantification methods. Next, we use a regression approach to calculate pairwise *similarities* among all IS (Figure 2B), which are represented as a similarity matrix *S*. The resulting similarities feed into the subsequent *clustering* algorithm that identifies co-occurrent IS by their abundance under the hypothesis that different IS generated by the same cell clones share the same quantification (Figure 2C). Since the real number of clones to be detected is unknown, we apply the similarity clustering algorithm for all sensible number of clusters k and select the best solution by comparing the cluster qualities (using a *silhouette score (Rousseeuw, 1987)* accounting for high inner-cluster similarity and low outer-cluster similarity). In a final step, *clonal abundances are reconstructed* by accounting for the identified multiplicity of integrations (Figure 2D). To quantify the similarity of two clusterings, especially when comparing a known ground truth with the results of the reconstruction pipeline, we use the adjusted Rand index (ARI) (Hubert & Arabie, 1985; Rand, 1971). Technical details are provided in the Supplementary Materials.

### Simulation data

In order to test and validate the clonal reconstruction pipeline within a scalable context, we generated *in silico* data based on a corresponding and published mathematical modeling approach to describe clonal tracking data (Baldow, Thielecke, & Glauche, 2016). It is the advantage of this simulation model that the true association of IS to clones (‘‘ground truth”) is intrinsically known. Technically, the time courses are generated from a stochastic, single cell-based model of clonal dynamics in which cells of different clones proliferate and differentiate with rates that are kept fixed for each clone. The differentiation rate for each clone *c* is initialized from a normal distribution 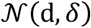, while the proliferation rate is dynamically regulated by a logistic growth function with overall carrying capacity *K* and maximal proliferation rate *p_max_* (which is identical for all clones, see Suppl. Table S4 for parameter values). Clone sizes 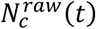 are given by the absolute number of cells belonging to a clone *c* at time point *t*.

For each clone *c* we initially assign a certain number of unique IS according to a Poisson distribution *Pois*(λ) in which λ reflects the average VCN for the particular transduction setting. The raw abundance *I^raw^* (i.e. without any measurement noise) of each individual IS *i* belonging to a clone *c* at time *t* is given as

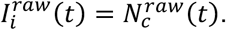

In order to account for the *measurement noise* of the detection process, we add a multiplicative noise 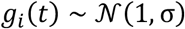 to the readout of every IS measurement

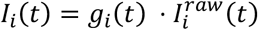

to obtain the time courses *I*(*t*) for all IS.

In order to mimic changing clonal contributions to different hematopoietic cell types, we introduce an additional inter-clonal shift, termed *clonal variability ν* that superimposes a random factor 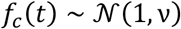 at each time point of measurement *t*, such that the abundance of simulated IS *I*\*(*t*) is given as

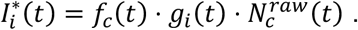

Herein, the first factor is clone specific and affects all IS of the same clone, while the second factor accounts for an individual measurement error for each IS. Following the above motivation, the temporal dimension *t* in the simulated data reflects both time *and* cell type in an *in vivo* study. The *relative* IS abundance is calculated as

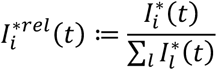

We varied the average VCN λ, the clonal variability ν, the measurement noise *σ*, as well as the number of measurements to generate a range of different model realizations for which the reconstruction process has been tested.

### Experimental data

#### Validation assay

We prepared a set of seven samples containing variable proportions of four different cell clones (see Supplementary Table S1) mixed into a background of transduced cells. Each of the four clones is characterized by a different number of IS (ID27 (1 IS), ID30 (4 IS), ID37 (6 IS), ID46 (10 IS)), while for the background only the average number of IS was determined to be approximately 1.8 copies. The prepared mixtures were primarily intended to address issues of sensitivity and reproducibility, while the variable abundance of four different clones serve as a suitable sample to verify whether the suggested bioinformatical pipeline correctly assigns the IS to their respective clones. For all samples, LV/genomic junctions were retrieved by SLiM PCR (see Supplementary Table S2), sequenced using NGS technologies, and mapped on the mouse genome to identify the nearest RefSeq gene. The relative amount of each clone in the mix was quantified and confirmed by droplet digital PCR (ddPCR) assay where primers and probes were specifically designed for at least one of the LV/genome junctions of each clone. As a filtering step, we are only considering IS that present 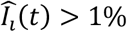 in any sample, thereby excluding IS from the background of other transduced cells.

#### Clonal tracing in mice

We used data from a set of mouse experiments in which wild type recipient mice were transplanted with *Cdkn2a^-/-^* BM-derived lineage-negative cells (lin-, HSPCs equivalent) carrying multiple vector integrations (average VCN *λ* ≈ 6 to 10). Lin-cells were transduced with the previously described self-inactivating lentiviral vector (SINLV) expressing GFP (SINLV.PGK.GFP) (Montini et al., 2009; Montini et al., 2006). Blood samples were taken at three to four different time points post transplantation and FACS sorted according to the phenotype of B-(CD19+), T-(CD3+), mononuclear, and myeloid (CD11b+) cells. Furthermore, at autopsy, from each mouse the bone marrow, blood, spleen, thymus, and lymph nodes were collected for IS site retrieval, providing the IS of the dominant/tumoral clone infiltrating the different tissues. In all these samples, LV/genomic junctions were retrieved from DNA samples by SLiM PCR (see Supplementary Materials), sequenced and mapped on the mouse genome to identify the nearest RefSeq gene. Unique IS within the tissue samples could be used to delineate the IS found in the pooled blood samples.

## Acknowledgments

We thank Thomas Zerjatke, Lars Thielecke and Friedemann Uschner for critical discussions of our manuscript.

This work was supported the German Federal Ministry of Research and Education (BMBF, grant number 031A315 “MessAge”) to I.G. and Telethon Foundation TGT16B01 and TGT16B03 to E.M. and Giovani Ricercatori Grant 2016 from the Italian Ministry of Health to A.C. and D.C. (GR-2016–02363681).

## Author contributions

S.W., C.B. and I.G. conceived and designed the study. S.W., C.B. developed computational tools and performed analysis. L.R. and P.G. performed the experiments under the supervision of D.C. and E.M.. A.C. and D.C. analyzed the experimental data. S.W., C.B., D.C., A.C. and I.G. wrote the paper. All authors reviewed the final manuscript.

## References

Aiuti, A., Biasco, L., Scaramuzza, S., Ferrua, F., Cicalese, M. P., Baricordi, C., … Naldini, L. (2013). Lentiviral hematopoietic stem cell gene therapy in patients with Wiskott-Aldrich syndrome. Science, 341(6148), 1233151. doi:10.1126/science.1233151

Baldow, C., Thielecke, L., & Glauche, I. (2016). Model Based Analysis of Clonal Developments Allows for Early Detection of Monoclonal Conversion and Leukemia. PLoS One, 11(10), e0165129. doi:10.1371/journal.pone.0165129

Beard, B. C., Adair, J. E., Trobridge, G. D., & Kiem, H. P. (2014). High-throughput genomic mapping of vector integration sites in gene therapy studies. Methods Mol Biol, 1185, 321–344. doi:10.1007/978-1-4939-1133-2_22

Berry, C. C., Gillet, N. a., Melamed, A., Gormley, N., Bangham, C. R. M., & Bushman, F. (2012). Estimating Abundances of Retroviral Insertion Sites from DNA Fragment Length Data. Bioinformatics (Oxford, England), 1–8. doi:10.1093/bioinformatics/bts004

Biffi, A., Montini, E., Lorioli, L., Cesani, M., Fumagalli, F., Plati, T., … Naldini, L. (2013). Lentiviral hematopoietic stem cell gene therapy benefits metachromatic leukodystrophy. Science (New York, N.Y.), 341(6148), 1233158–1233158. doi:10.1126/science.1233158

Cavazzana, M., Bushman, F. D., Miccio, A., Andre-Schmutz, I., & Six, E. (2019). Gene therapy targeting haematopoietic stem cells for inherited diseases: progress and challenges. Nat Rev Drug Discov, 18(6), 447–462. doi:10.1038/s41573-019-0020-9

Cavazzana-Calvo, M., Payen, E., Negre, O., Wang, G., Hehir, K., Fusil, F., … Leboulch, P. (2010). Transfusion independence and HMGA2 activation after gene therapy of human β-thalassaemia. Nature, 467(7313), 318–322. doi:10.1038/nature09328

Cesana, D., Ranzani, M., Volpin, M., Bartholomae, C., Duros, C., Artus, A., … Montini, E. (2014). Uncovering and Dissecting the Genotoxicity of Self-inactivating Lentiviral Vectors In Vivo. Molecular therapy: the journal of the American Society of Gene Therapy, 22(4), 774–785. doi:10.1038/mt.2014.3

Ferrari, G., Thrasher, A. J., & Aiuti, A. (2021). Gene therapy using haematopoietic stem and progenitor cells. Nat Rev Genet, 22(4), 216–234. doi:10.1038/s41576-020-00298-5

Firouzi, S., López, Y., Suzuki, Y., Nakai, K., Sugano, S., Yamochi, T., & Watanabe, T. (2014). Development and validation of a new high-throughput method to investigate the clonality of HTLV-1-infected cells based on provirus integration sites. Genome Medicine, 6(6), 46. doi:10.1186/gm568

Hacein-Bey Abina, S., Gaspar, H. B., Blondeau, J., Caccavelli, L., Charrier, S., Buckland, K., … Cavazzana, M. (2015). Outcomes Following Gene Therapy in Patients With Severe Wiskott-Aldrich Syndrome. Jama, 313(15), 1550–1563. doi:10.1001/jama.2015.3253

Hacein-Bey-Abina, S., von Kalle, C., Schmidt, M., Le Deist, F., Wulffraat, N., McIntyre, E., … Fischer, A. (2003). A serious adverse event after successful gene therapy for X-linked severe combined immunodeficiency. N Engl J Med, 348(3), 255–256. doi:10.1056/NEJM200301163480314

Hubert, L., & Arabie, P. (1985). Comparing partitions. Journal of Classification, 2(1), 193–218. doi:10.1007/BF01908075

Kohn, D. B., Booth, C., Kang, E. M., Pai, S. Y., Shaw, K. L., Santilli, G., … consortium, N. C. (2020). Lentiviral gene therapy for X-linked chronic granulomatous disease. Nat Med. doi:10.1038/s41591-019-0735-5

Marktel, S., Scaramuzza, S., Cicalese, M. P., Giglio, F., Galimberti, S., Lidonnici, M. R., … Ferrari, G. (2019). Intrabone hematopoietic stem cell gene therapy for adult and pediatric patients affected by transfusion-dependent ß-thalassemia. Nat Med, 25(2), 234–241. doi:10.1038/s41591-018-0301-6

Montini, E., Cesana, D., Schmidt, M., Sanvito, F., Bartholomae, C. C., Ranzani, M., … Naldini, L. (2009). The genotoxic potential of retroviral vectors is strongly modulated by vector design and integration site selection in a mouse model of HSC gene therapy. J Clin Invest, 119(4), 964–975. doi:10.1172/JCI37630

Montini, E., Cesana, D., Schmidt, M., Sanvito, F., Ponzoni, M., Bartholomae, C., … Naldini, L. (2006). Hematopoietic stem cell gene transfer in a tumor-prone mouse model uncovers low genotoxicity of lentiviral vector integration. Nat Biotechnol, 24(6), 687–696. doi:10.1038/nbt1216

Notta, F., Zandi, S., Takayama, N., Dobson, S., Gan, O. I., Wilson, G., … Dick, J. E. (2016). Distinct routes of lineage development reshape the human blood hierarchy across ontogeny. Science, 351(6269), aab2116–aab2116. doi:10.1126/science.aab2116

Rand, W. M. (1971). Objective Criteria for the Evaluation of Clustering Methods. Journal of the American Statistical Association, 66(336), 846–850. doi:10.1080/01621459.1971.10482356

Rousseeuw, P. J. (1987). Silhouettes: A graphical aid to the interpretation and validation of cluster analysis. Journal of Computational and Applied Mathematics, 20, 53–65. doi:10.1016/0377-0427(87)90125-7

Scala, S., & Aiuti, A. (2019). In vivo dynamics of human hematopoietic stem cells: novel concepts and future directions. Blood Adv, 3(12), 1916–1924. doi:10.1182/bloodadvances.2019000039

Scala, S., Basso-Ricci, L., Dionisio, F., Pellin, D., Giannelli, S., Salerio, F. A., … Biasco, L. (2018). Dynamics of genetically engineered hematopoietic stem and progenitor cells after autologous transplantation in humans. Nat Med, 24(11), 1683–1690. doi:10.1038/s41591-018-0195-3

Sessa, M., Lorioli, L., Fumagalli, F., Acquati, S., Redaelli, D., Baldoli, C., … Biffi, A. (2016). Lentiviral haemopoietic stem-cell gene therapy in early-onset metachromatic leukodystrophy: an ad-hoc analysis of a non-randomised, open-label, phase 1/2 trial. The Lancet. doi:10.1016/s0140-6736(16)30374-9

