## Supplementary Materials for "Clonal reconstruction from co-occurrence of vector integration sites allows accurate quantification of expanding clones in vivo"

for

### Technical description of the reconstruction pipeline

Similarity of IS abundances

We applied an $R^2$ approach to quantify the similarity in the abundance of two IS, which is based on the *relative* IS reads $I_i^{*rel}(t) := \frac{I_i(t)}{\sum_l I_l(t)}$ , either obtained from simulated or measured data. Comparing the $I_i^{*rel}(t)$ for all possible pairs of filtered IS *i, j*, and calculating a linear regression through the origin (compare Figure 2B) we obtain the coefficient of determination $R_{i,j}^2$ which is interpreted as a normalized measure of how similar the two measurement series are (with $R^2 \rightarrow 1$ indicating perfect correlation). Based hereon, we construct a similarity matrix *S* that contains the calculated values for all pairs of IS, i.e. $S_{i,j} = R_{i,j}^2$. Prior to clustering, we obtain a rescaled similarity matrix *S'*, in which similarity values are linearly rescaled such that the lowest value is set to 0 and the largest value is set to 1.

We point out that the regression approach based on $R^2$ can be replaced by alternative methods for the pairwise evaluation of IS similarity, such as a Canberra distance (Lance & Williams, 1966).

Clustering of IS

Clustering algorithms are a suitable tool to identify sets of IS with correlated behaviors, which indicate a common clonal origin. Technically, each IS *i* within the rescaled similarity matrix *S'* is assigned to one cluster *c* out of a number of available clusters *k*. Clustering algorithms aim to maximize the inner-cluster similarities and minimize outer-cluster similarities to obtain an optimal assignment of the elements to the *k* clusters. As the number of expected clusters (i.e. clones, to which the IS are assigned) is unknown *a priori*, we apply the cluster algorithm multiple times (compare Figure 2D) while varying the number of clusters *k* between 2 and the number of filtered IS minus 1 (the later representing the case that each except for one IS would belong to an individual clone; the upper bound was chosen as some clustering algorithms do not support assigning each element to its own cluster, i.e. *k* = |*IS*|). We use *partitioning around medoids* (PAM) as a suitable clustering algorithm, which is a realization of the k-medoids algorithm (Kaufman & Rousseeuw, 1990). In contrast to a k-means approach, PAM does not choose an artificial center for the clusters, but a single, central data point in the cluster. Also, instead of minimizing the distances to the centers, PAM minimizes the sum of all pairwise distances within each cluster. Compared to k-means, PAM is regarded to be more robust to outliers.

Evaluation of clusters

In order to compare clustering results for different numbers of clusters $k$ we use the *silhouette score* (Rousseeuw, 1987). This score quantifies the quality of a given clustering by comparing the inner-cluster similarity with the outer-cluster similarity.

Technically, a silhouette value $s$ of IS $i$ is defined as:

$$sil(i) = \frac{oc(i) - ic(i)}{max\{ic(i), oc(i)\}}$$

Here *ic(i)* and *oc(i)* are the average similarities of $i$ to all IS within the same and the IS within other clusters, respectively. The overall silhouette score for a given number of clusters $k$ is defined as the average of all individual silhouette values over all IS:

$$\overline{sil}(k) = \frac{\sum_{i=1}^{|IS|} s(i)}{|IS|}$$

A high silhouette score indicates a strong inner-cluster similarity compared to a weak outer-cluster similarity. Calculating the corresponding silhouette scores $\overline{sil}(k)$ for different numbers of clusters $2 < k < |IS| - 1$, we consider the maximum $k$, named $k^*$, to identify the most likely number of clusters for the selected IS.

Calculation of relative clonal abundances

According to the optimal clustering, all IS are assigned to one of $k^*$ clusters, each of which is interpreted as an independent clone. As a consequence, the clonal abundance $\widetilde{N_c}(t)$ is now given as the average over all IS abundances $\widehat{I_i}(t)$ assigned to the same cluster $c$. Additionally, all unfiltered IS abundances (that will remain un-clustered to their source clones) have to be rescaled with an average number of IS per clone (representing an approximation of the average VCN) to correct for their potential co-occurrence within the same clone. Although this approach does not allow to interpret unfiltered IS as separate clones, it corrects their relative abundance to the observed clusters.

Comparison of the reconstruction results with a known ground truth

In order to evaluate the reconstruction quality, we compare the results of the clonal reconstruction process to the known ground truth that is available from simulated time courses and selected, annotated experiments. Technically, we calculate the adjusted Rand index (ARI) as a measure of overlap between the ground truth association and the clustering result. A value of ARI = 1 indicates a perfect clustering, i.e. IS that were associated to the same clone in the ground truth are also jointly recovered while other IS remain in distinct clusters. An ARI value of 0 would indicate a completely random assignment of IS to clusters. As the ARI only uses information about the co-occurrence of elements, it is label-independent.

**Experimental data**

Simulation model

The simulation model is adapted from (Baldow, Thielecke, & Glauche, 2016), in which time courses are generated from a stochastic, single cell-based model of clonal dynamics. The differentiation rate for each clone $c$ is initialized from a normal distribution $\mathcal{N}(\mathrm{d}, \mathrm{s_d})$ and kept fixed thereafter. The proliferation rate is identical for all clones and regulated by a logistic growth function with overall carrying capacity $K$ and maximal proliferation rate $p_{max}$. The chosen model parameters are provided in Suppl. Table S4.

Validation assay.

Chronic Myelogenous Leukemia (K562) cell clones were transduced with a lentiviral vector expressing GFP under the control of the Spleen Focus Forming Virus promoter (Montini et al., 2009), using a multiplicity of infection (MOI) of 10. Two weeks after transduction, single cell clones were isolated from the GFP+ population of cells by FACS sorting and cultured in IMDM-medium for several weeks. For each clone, the LV vector/genome junctions were retrieved by sonication-based linker-mediated (SLiM) PCR. In Supplementary Table S2 we reported the different IS identified in each of the four clones used in the validation assay. Additionally, human B-lymphoblastoid cells (JY cells) were transduced in bulk with a lentiviral vector expressing GFP under the control of PGK promoter (Montini et al., 2006) and using a multiplicity of infection of 1. The VCN in the K562 cell clones and JY was determined by ddPCR.

Clonal tracing in mice.

In agreement with previously published data (Montini et al., 2006), mice receiving Cdkn2a$^{-/-}$ Lin- cells transduced with SINLV.PGK.GFP developed tumors similarly to the mock-control group.

To recover enough starting material for the sorting procedure, DNA extraction and subsequent molecular analyses, an equal amount of blood from a cohort of two to three different animals was pooled together prior to sorting. The composition of each pool was maintained constant throughout the whole experiment, so that each pool is composed by the same mice over time.

Retrieval of IS from cell DNA / SLiM PCR

For the retrieval of vector IS from genomic DNA, we adopted a SLiM-PCR method similar to the previously described (Firouzi et al., 2014). Briefly, genomic DNA was sheared using a Covaris E220 Ultrasonicator (Covaris Inc., Woburn ,MA.), and split in three technical replicates. The fragmented DNA was then subjected to end repair and 3' adenylation using the NEBNext® Ultra™ DNA Library Prep Kit for Illumina® (New England Biolabs, Ipswich, MA.), and then ligated (DNA Technologies ligation kit, Skokie, IL.) to the linker cassettes (LC) containing a sequence barcode for sample identification and all the sequences required for the Read 2 Illumina paired end sequencing. Ligation products were then subjected to 35 cycles of exponential PCR and next ten additional PCR cycles were done to add sequences required for sequencing. Finally, the amplification products were sequenced using the Illumina NextSeq 550 sequencing platform (Illumina, San Diego, CA.).

Sequencing reads were then processed by a dedicated bioinformatics pipeline (VISPA2) as described before (Spinozzi et al., 2017). Briefly, paired sequence reads are filtered for quality standards, barcodes identified for sample de-multiplexing of the sequence reads, the cellular genomic sequence mapped on the reference Human genome (Human Genome_GRCh37/hg19 Feb. 2019) or mouse (Mouse Genome_mm9) and the nearest RefSeq gene assigned to each unambiguously mapped integration site. For the quantification of the abundance of each clone we adopted an estimation method previously described (Berry et al., 2012) where the abundance is determined by the number of different DNA fragments containing the same vector/cell genome junctions flanked by a genomic segment variable in size depending on the shear site position and that will be unique for each different cell genome present in the starting cell population. Therefore, the number of different shear sites assigned to an IS will be proportional to the initial number of contributing cells, allowing to estimate the clonal abundance in the starting sample avoiding the biases introduced by PCR amplification.

Droplet digital (dd)PCR

The abundance of each K562 clone in the DNA mixes was measured by ddPCR using the QuantaLife ddPCR system. Custom-made ddPCR assays were designed to specifically amplify at least one of the LV vector genome junctions retrieved in each clone. Human GAPDH ddPCR assay was used as the housekeeping gene. 20-30 nanograms of genomic DNA for each mix were used for PCR amplification performed in triplicate and in a final volume of 20 μl. Plates were quantified in a QuantaLife droplet reader, and the concentrations of the targets in the samples were determined using QuantaSoft software.

**R package MultIS**

The R package *MultIS* provides an implementation of the functionality presented in the accompanying manuscript. The package is available from the Comprehensive R Archive Network (CRAN) under the Lesser GNU Public License Version 3 (LGPLv3).

The package comprises the basic methods used for filtering, calculation of similarities, clustering, and the evaluation of clusters. Furthermore, it provides corresponding visualization routines for each step in the pipeline. Along with the package we provide a vignette illustrating an exemplary workflow.

Methods for normalizing and filtering IS read data

*MultIS* implements functionality for the normalization and filtering of time course data with an intuitive naming of the respective functions.

The first recommended step to achieve comparability within one experimental series is a normalization of IS counts to a relative scale for each measurement. For the subsequent filtering step, different methods can be adapted, e.g. with respect to an absolute number of selected, most abundant IS or with respect to a minimal threshold.

The filtering methods also include matching for patterns in measurements or in IS names. *MultIS* further includes different functions to transform measurement annotations and shorten IS identifiers to their shortest distinct prefix.

Measuring similarities between IS

To measure the similarity between IS we construct a similarity matrix $S$ that contains each pairwise rating of similarity. We recommend using a $R^2$ metric, while other metrics can be considered as well. Technically, *MultIS* additionally supports all other methods provided by "stats::dist" package such as the Euclidean, Manhattan, or Canberra (Lance & Williams, 1966) distance.

Clustering IS

We use partitioning around medoids (PAM) (Kaufman & Rousseeuw, 1990) as a suitable method to identify clusters of IS with correlated behaviors. However, the cluster function can be called with other arguments than "kmedoids" (which is an implementation of PAM), such as "kmeans" (MacQueen, 1967; Steinhaus, 1956), or any method implemented in "stats::hclust".

Rating and finding the optimal number of clusters

In the experimental setting the number of clones in a sample is unknown. Thus, our method aims towards estimating this number from the correlation structure within the rescaled similarity matrix $S'$. Supervised clustering methods require a predefined number of expected clusters k. We repeatedly apply the clustering for all values of $k$ between 2 and the number of IS – 1. We suggest to use the silhouette score (Rousseeuw, 1987) to compare the cluster results for different given values of *k* and to identify the most optimal number k* that maximizes the inner and minimizes the outer-cluster similarity. Alternative methods can be used instead, e.g. the SD index (Halkidi, Vazirgiannis, & Batistakis, 2000), the point-biserial index (Kraemer, 2006; Milligan, 1980, 1981), or the Dunn index (Dunn†, 2008).

<u>Methods for the visualization of the steps in our pipeline</u>
Each step in the reconstruction pipeline of "*MultIS*" returns an object that can directly be used with R's "plot" function to visualize the results and to give the user a quick overview of the data.

*Time courses*
Time courses are visualized using a stacked area plot that shows their relative contribution over time. The measurements can either be time points or combinations of time points and cell types. If the cell type is encoded in the measurement name, we provide the functionality to extract the cell type part from the naming convention of the measurement and use this as facets of different sub-plots. Coloring schemes can be applied to ensure an internal consistency across different plots.

*Similarity matrices*
Similarity matrices *S* can be efficiently visualized as heatmaps. Internally, the heat map function also performs a hierarchical clustering to display similar IS adjacent to each other.

*Clustered IS and similarities*
We recommend a spring model to visualize the results of different clustering methods. IS are laid out as nodes on a plane and connected via springs. The strength of these springs is proportional to the similarity contained in the similarity matrix *S*. Thus, more closely related IS take a position that is closer to each other in this plot. The shading of the edges refers to the mutual similarity of any two IS. The coloring of the nodes (i.e. IS) indicates whether they are associated with the same cluster. If an additional ground truth is known, an inner and outer coloring of the nodes can be used to distinguish the reconstructed and the known associations.

### Data provision and reproducibility

We provide an R-script file along with all the necessary data sets to fully reproduce all steps within our manuscript (https://gitlab.com/imb-dev/clonal-reconstruction-figures). The script relies upon the functionality of the "*MultIS*" package, outlined above.

### References


Baldow, C., Thielecke, L., & Glauche, I. (2016). Model Based Analysis of Clonal Developments Allows for Early Detection of Monoclonal Conversion and Leukemia. *PLoS One, 11*(10), e0165129. doi:10.1371/journal.pone.0165129

Berry, C. C., Gillet, N. a., Melamed, A., Gormley, N., Bangham, C. R. M., & Bushman, F. (2012). Estimating Abundances of Retroviral Insertion Sites from DNA Fragment Length Data. *Bioinformatics (Oxford, England)*, 1-8. doi:10.1093/bioinformatics/bts004

Dunn†, J. C. (2008). Well-Separated Clusters and Optimal Fuzzy Partitions. *Journal of Cybernetics, 4*(1), 95-104. doi:10.1080/01969727408546059

Firouzi, S., López, Y., Suzuki, Y., Nakai, K., Sugano, S., Yamochi, T., & Watanabe, T. (2014). Development and validation of a new high-throughput method to investigate the clonality of HTLV-1-infected cells based on provirus integration sites. *Genome Medicine, 6*(6), 46. doi:10.1186/gm568

Halkidi, M., Vazirgiannis, M., & Batistakis, Y. (2000). *Quality Scheme Assessment in the Clustering Process*, Berlin, Heidelberg.

Kaufman, L., & Rousseeuw, P. J. (1990). Partitioning Around Medoids (Program PAM). In *Finding Groups in Data* (pp. 68-125).

Kraemer, H. (2006). Biserial Correlation. *1*, 276-279. doi:10.1002/0471667196.ess0153.pub2

Lance, G. N., & Williams, W. T. (1966). Computer Programs for Hierarchical Polythetic Classification ("Similarity Analyses"). *The Computer Journal, 9*(1), 60-64. doi:10.1093/comjnl/9.1.60

MacQueen, J. (1967). *Some methods for classification and analysis of multivariate observations.* Paper presented at the Proceedings of the fifth Berkeley symposium on mathematical statistics and probability.

Milligan, G. W. (1980). An examination of the effect of six types of error perturbation on fifteen clustering algorithms. *Psychometrika, 45*(3), 325-342. doi:10.1007/bf02293907

Milligan, G. W. (1981). A monte carlo study of thirty internal criterion measures for cluster analysis. *Psychometrika, 46*(2), 187-199. doi:10.1007/bf02293899

Montini, E., Cesana, D., Schmidt, M., Sanvito, F., Bartholomae, C. C., Ranzani, M., . . . Naldini, L. (2009). The genotoxic potential of retroviral vectors is strongly modulated by vector design and integration site selection in a mouse model of HSC gene therapy. *J Clin Invest, 119*(4), 964-975. doi:10.1172/JCI37630

Montini, E., Cesana, D., Schmidt, M., Sanvito, F., Ponzoni, M., Bartholomae, C., . . . Naldini, L. (2006). Hematopoietic stem cell gene transfer in a tumor-prone mouse model uncovers low genotoxicity of lentiviral vector integration. *Nat Biotechnol, 24*(6), 687-696. doi:10.1038/nbt1216

Rousseeuw, P. J. (1987). Silhouettes: A graphical aid to the interpretation and validation of cluster analysis. *Journal of Computational and Applied Mathematics, 20*, 53-65. doi:10.1016/0377-0427(87)90125-7

Spinozzi, G., Calabria, A., Brasca, S., Beretta, S., Merelli, I., Milanesi, L., & Montini, E. (2017). VISPA2: a scalable pipeline for high-throughput identification and annotation of vector integration sites. *BMC Bioinformatics, 18*(1). doi:10.1186/s12859-017-1937-9

Steinhaus, H. (1956). Sur la division des corps matériels en parties. *Bull. Acad. Polon. Sci, 1*(804), 801.
