## Supplementary Figures for "Clonal reconstruction from co-occurrence of vector integration sites allows accurate quantification of expanding clones in vivo"

Supplementary Figure S1

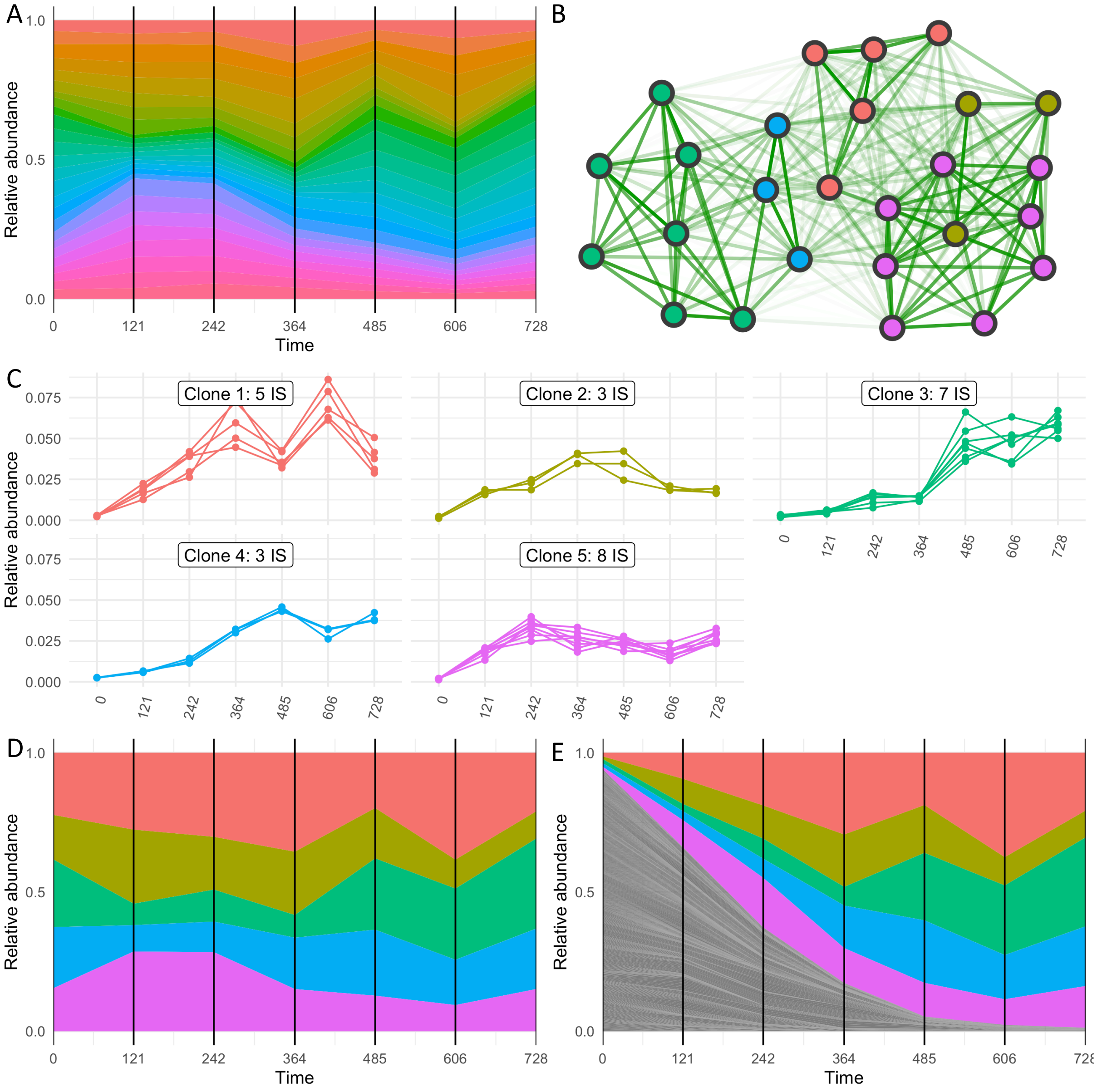


**Figure S1 Reconstruction for a simulated time course with low measurement noise σ = 0.15.**

Time courses are initialized with 100 clones, each with 100 identical cells. The number of IS per clone are chosen from a Poisson distribution with average VCN λ=5. **A** relative abundances of IS as a function of time after applying the filtering step. **B** The similarity (indicated by edge brightness) between each pair of integration sites is superimposed by the optimal clustering (indicated by color of the nodes) obtained from the reconstruction pipeline. **C** time series of all IS assigned to the same clusters/clones (color coding corresponds to subfigure B). **D** corrected clonal time series for the five identified clones. **E** shows the corrected clonal time course together with the IS that did not pass the initial filtering step (grey).

Supplementary Figure S2

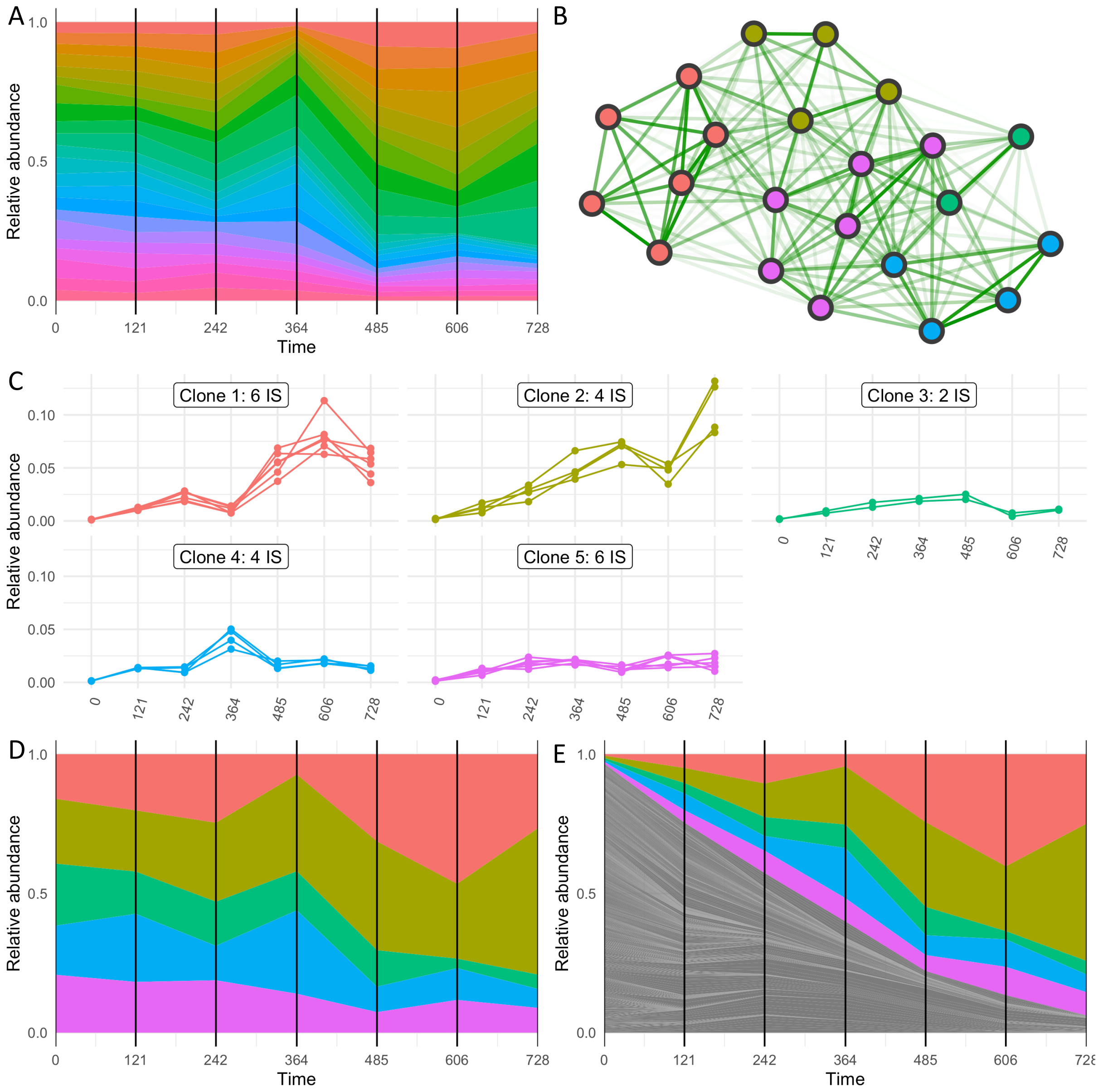


**Figure S2 Reconstruction for a simulated time course with high measurement noise σ = 0.25.**

Time courses are initialized with 100 clones, each with 100 identical cells. The number of IS per clone are chosen from a Poisson distribution with average VCN $\lambda$=5. **A** relative abundances of IS as a function of time after applying the filtering step. **B** The similarity (indicated by edge brightness) between each pair of integration sites is superimposed by the optimal clustering (indicated by color of the nodes) obtained from the reconstruction pipeline. **C** time series of all IS assigned to the same clusters/clones (color coding corresponds to subfigure B). **D** corrected clonal time series for the five identified clones. **E** shows the corrected clonal time course together with the IS that did not pass the initial filtering step (grey).

Supplementary Figure S3

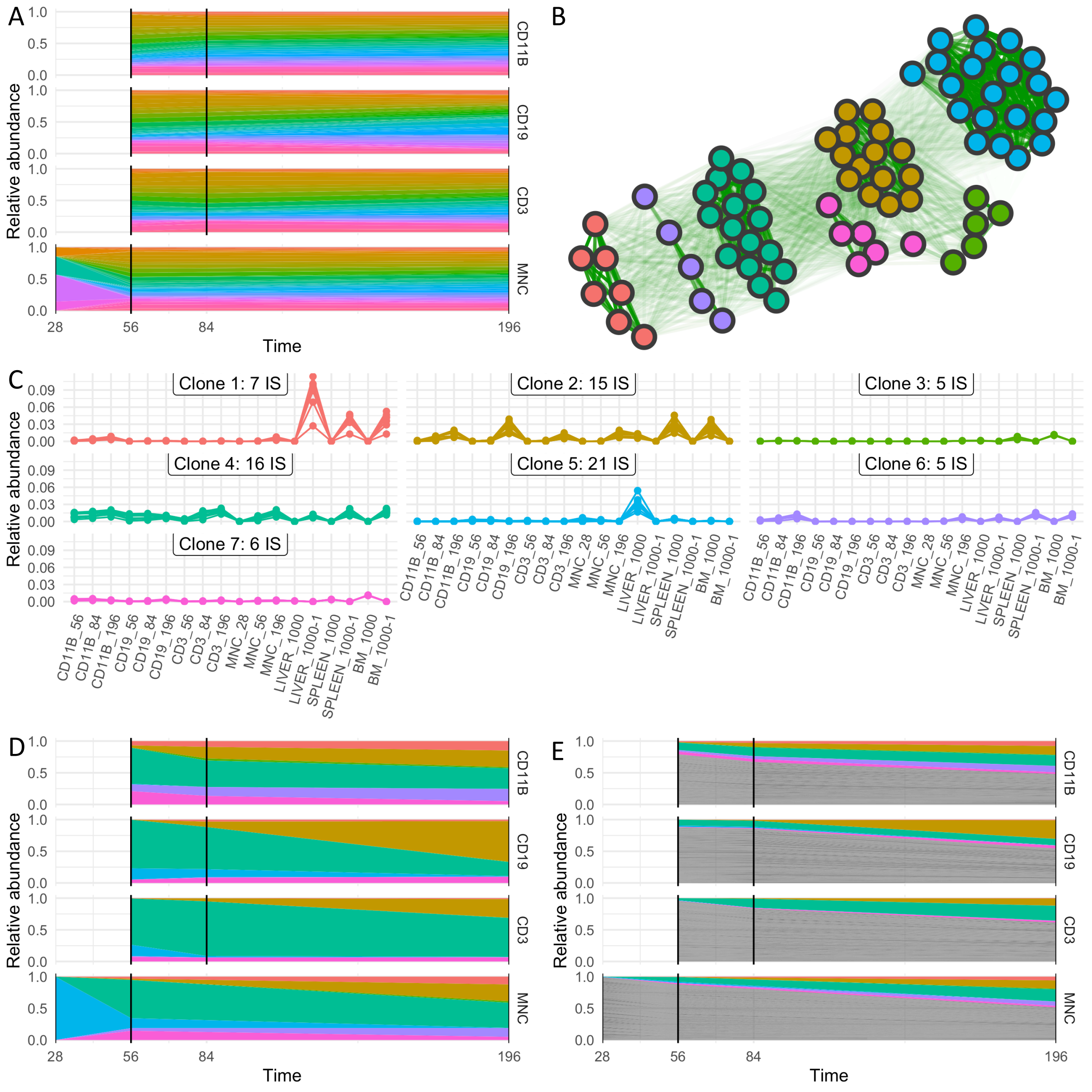


**Figure S3 Experimental data and clonal reconstruction for wild type mouse E3D.**

Mice were transplanted with HSPCs previously transduced with a PGK-based vector. Collected cells were sorted according to their immune phenotype and analyzed separately for IS abundance.

**A** relative abundances of IS as a function of time for CD11b, CD19, CD3, and mononuclear cells, for which multiple measurements are available. **B** similarity (indicated by edge brightness) between each pair of integration sites superimposed by the optimal clustering (indicated by color of the nodes) obtained from the reconstruction pipeline. **C** time series of all IS assigned to the same clusters/clones (color coding corresponds to subfigure B). **D** shows the corrected clonal time course for the seven identified clones. **E** shows the corrected clonal time series together with the IS that did not pass the initial filtering step (grey).

Supplementary Figure S4

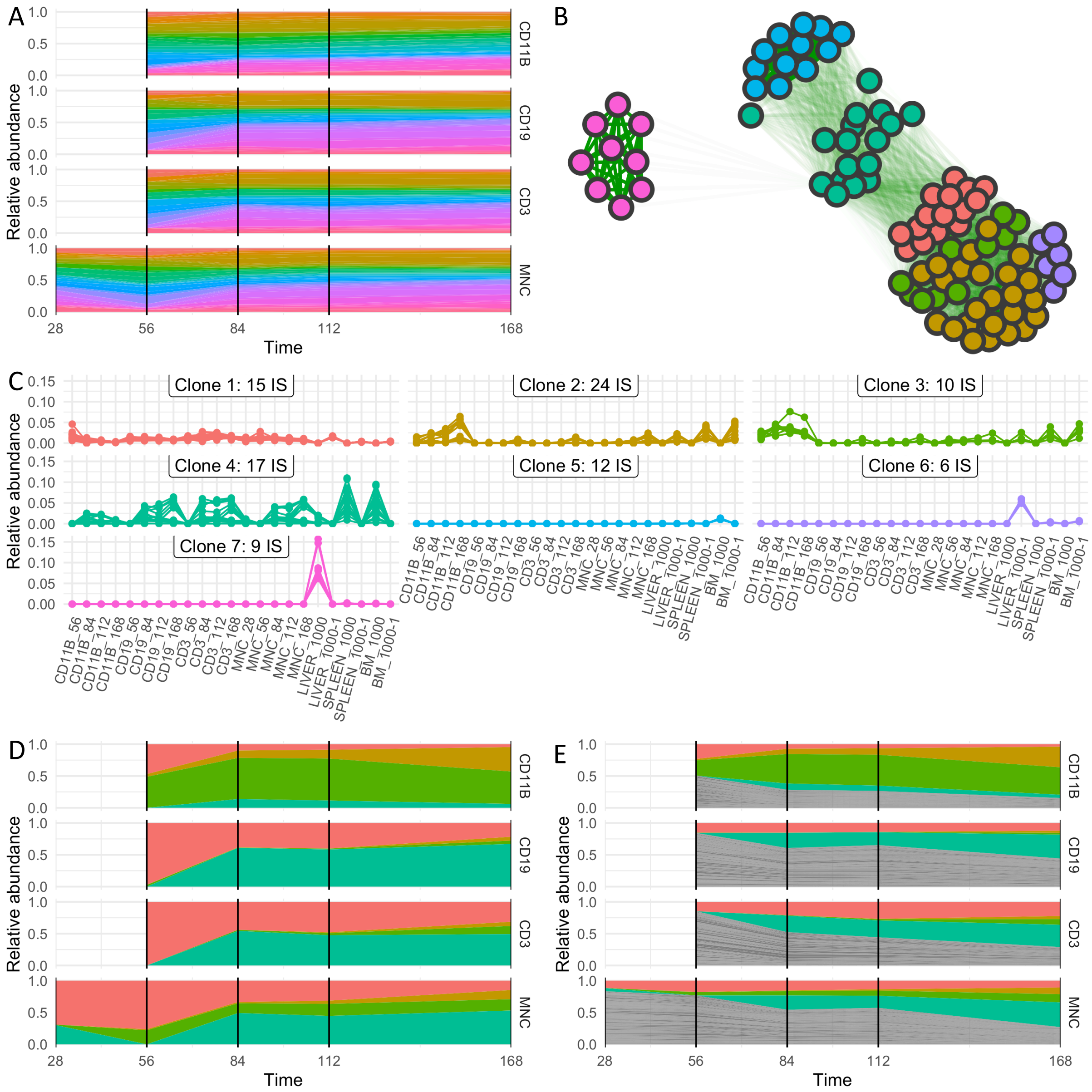


**Figure S4 Experimental data and clonal reconstruction for wild type mouse E2L.**

Mice were transplanted with HSPCs previously transduced with a PGK-based vector. Collected cells were sorted according to their immune phenotype and analyzed separately for IS abundance.

**A** relative abundances of IS as a function of time for CD11b, CD19, CD3, and mononuclear cells, for which multiple measurements are available. **B** similarity (indicated by edge brightness) between each pair of integration sites superimposed by the optimal clustering (indicated by color of the nodes) obtained from the reconstruction pipeline. **C** time series of all IS assigned to the same clusters/clones (color coding corresponds to subfigure B). **D** shows the corrected clonal time course for the seven identified clones. **E** shows the corrected clonal time series together with the IS that did not pass the initial filtering step (grey).

**Table S1** Relative composition of the different mixes for the validation assay. JY indicating a polyclonal background.

| Volume | Mix 1 | Mix 2 | Mix 3 | Mix 4 | Mix 5 | Mix 6 | Mix 7 |
|---|---|---|---|---|---|---|---|
| JY | 0.60 | 0.35 | 0.10 | 0.09 | 0.07 | 0.24 | 0.56 |
| ID#27 | 0.05 | 0.05 | 0.05 | 0.40 | 0.03 | 0.08 | 0.25 |
| ID#30 | 0.25 | 0.50 | 0.75 | 0.03 | 0.40 | 0.25 | 0.08 |
| ID#37 | 0.05 | 0.05 | 0.05 | 0.08 | 0.25 | 0.40 | 0.03 |
| ID#46 | 0.05 | 0.05 | 0.05 | 0.40 | 0.25 | 0.03 | 0.08 |

**Table S2** Genomic position of clone specific IS for the K562 cell clones

| Clone ID(a) | N IS(b) | Chr(c) | Integration_locus(d) | Gene Name (e) |
|---|---|---|---|---|
| ID#27 | 1 | 5 | 72866480 | UTP15 |
| ID#30 | 4 | 2 | 223461674 | FARSB |
| | | 14 | 97342533 | VRK1 |
| | | 2 | 208325826 | CREB1 |
| | | 4 | 129335149 | LOC100507487 |
| ID#37 | 6 | 8 | 91674211 | TMEM64 |
| | | 18 | 38162304 | LINC01477 |
| | | 14 | 81652945 | GTF2A1 |
| | | 12 | 96398298 | LTA4H |
| | | X | 120619080 | MIR3672 |
| | | 7 | 110789918 | IMMP2L |
| ID#46 | 10 | 6 | 45172433 | SUPT3H |
| | | 19 | 8002970 | TIMM44 |
| | | 4 | 909971 | GAK |
| | | 11 | 8711542 | RPL27A |
| | | 1 | 160414727 | VANGL2 |
| | | 4 | 120428505 | PDE5A |
| | | 4 | 63782063 | ADGRL3-AS1 |
| | | 15 | 78770928 | IREB2 |
| | | 20 | 33384442 | NCOA6 |
| | | 6 | 47118860 | TNFRSF21 |

(a) **Clone ID:** identifier of the K562 clone
(b) **N IS:** number of different LV integrations identified
(c) **Chr:** chromosome number of the targeted gene in the human genome
(d) **Integration locus:** genomic coordinate of the integration point
(e) **Gene Symbol:** gene symbol of the targeted gene

**Table S3**: Expected and reconstructed clonal abundances for the validation assay (compare Figure 5). Red numbers indicate divergence between reconstructed and expected.

| | | Mix 1 | Mix 2 | Mix 3 | Mix 4 | Mix 5 | Mix 6 | Mix 7 |
|---|---|---|---|---|---|---|---|---|
| **Expectation** (rel. abundance without background) | ID#27 | 0.12 | 0.08 | 0.06 | 0.44 | 0.03 | 0.11 | 0.57 |
| | ID#30 | 0.62 | 0.77 | 0.83 | 0.03 | 0.43 | 0.33 | 0.18 |
| | ID#37 | 0.12 | 0.08 | 0.06 | 0.09 | 0.27 | 0.53 | 0.07 |
| | ID#46 | 0.12 | 0.08 | 0.06 | 0.44 | 0.27 | 0.04 | 0.18 |
| **Reconstruction** (naïve, ID#27 wrongly assigned to ID#46) | ~~ID#27~~ | 0.00 (-0.12) | 0.00 (-0.08) | 0.00 (-0.06) | 0.00 (-0.44) | 0.00 (-0.03) | 0.00 (-0.11) | 0.00 (-0.57) |
| | ID#30 | 0.74 (+0.12) | 0.86 (+0.09) | 0.90 (+0.06) | 0.07 (+0.03) | 0.46 (+0.03) | 0.36 (+0.03) | 0.39 (+0.20) |
| | ID#37 | 0.12 (-0.00) | 0.07 (-0.01) | 0.05 (-0.00) | 0.14 (+0.05) | 0.27 (-0.00) | 0.55 (+0.03) | 0.12 (+0.05) |
| | ID#46 | 0.14 (+0.01) | 0.07 (-0.00) | 0.05 (-0.00) | 0.79 (+0.36) | 0.27 (+0.00) | 0.09 (+0.05) | 0.49 (+0.31) |
| **Reconstruction** (corrected, reconstruction of four clones) | ID#27 | 0.15 (+0.03) | 0.10 (+0.02) | 0.07 (+0.01) | 0.53 (+0.09) | 0.04 (+0.01) | 0.14 (+0.03) | 0.64 (+0.07) |
| | ID#30 | 0.63 (+0.00) | 0.78 (+0.01) | 0.84 (+0.00) | 0.03 (-0.00) | 0.43 (+0.00) | 0.31 (-0.01) | 0.16 (-0.03) |
| | ID#37 | 0.10 (-0.02) | 0.06 (-0.01) | 0.05 (-0.01) | 0.07 (-0.02) | 0.25 (-0.02) | 0.48 (-0.05) | 0.05 (-0.02) |
| | ID#46 | 0.11 (-0.01) | 0.06 (-0.01) | 0.05 (-0.01) | 0.37 (-0.07) | 0.27 (+0.00) | 0.07 (+0.03) | 0.16 (-0.03) |

**Table S4**: Simulation Parameters

| | Figure 3 | Figure 4 | Figure S1 | Figure S2 |
|---|---|---|---|---|
| number of clones | 100 | 100 | 100 | 100 |
| cells per clone | 100 | 100 | 100 | 100 |
| number of measurements | 3 ... 15 | 3 ... 15 | 7 | 7 |
| VCN | 5 | 2, 5, 10 | 5 | 5 |
| clonal variability $\nu$ | 0 | 0, 0.125, 0.25, 0.375, 0.5 | 0.25 | 0.25 |
| measurement noise $\sigma$ | 0.025 | 0.025, 0.04 0.08 | 0.15 | 0.25 |
| proliferation rate $p_{max}$ | 0.06 | 0.06 | 0.06 | 0.06 |
| carrying capacity K | 10,000 | 10,000 | 10,000 | 10,000 |
| differentiation rate d | 0.04 | 0.04 | 0.04 | 0.04 |
| standard deviation of the differentiation rate $\delta$ | 0.0, 0.0025, 0.005 | 0.0025 | 0.01 | 0.01 |
